# Epigenetic profiling and incidence of disrupted development point to gastrulation as aging ground zero in *Xenopus laevis*

**DOI:** 10.1101/2022.08.02.502559

**Authors:** Bohan Zhang, Andrei E. Tarkhov, Wil Ratzan, Kejun Ying, Mahdi Moqri, Jesse R. Poganik, Benjamin Barre, Alexandre Trapp, Joseph A. Zoller, Amin Haghani, Steve Horvath, Leonid Peshkin, Vadim N. Gladyshev

## Abstract

Recent studies suggest the existence of a natural rejuvenation event during early embryonic development of mice, followed by epigenetic aging. Here, by profiling embryonic DNA methylation in the African clawed frog, *Xenopus laevis*, we found that the epigenetic entropy basepoint maps to the gastrulation stage of embryogenesis and corresponds to a rapid increase in embryo transcript abundance. We further developed a frog aging clock, revealing that this species epigenetically ages. Application of this clock to developmental stages identified a decrease in epigenetic age during early embryogenesis, with the minimal age reached around gastrulation. By examining individual developmental trajectories of 6,457 embryos, we found that this stage is also accompanied by a higher incidence of disrupted development. Taken together, our data point to gastrulation as a critical stage for aging and natural rejuvenation, characterized by the lowest epigenetic age, increased mortality, nadir of DNA methylation entropy and rapid increase in embryo transcript abundance, defining aging ground zero as the basepoint where rejuvenation ends and the aging process begins.

## Introduction

Recent experimental evidence begins to change the conventional notion that aging is irreversible^1^. A series of experiments involving cell reprogramming has shown that adult cells can transition to an embryonic-like state where their biological age is lower than their original biological age following the induction of reprogramming by several key transcription factors or treatment with chemicals^2–9^. This process, also known as rejuvenation, suggests the possibility of biological age reversal of cells and tissues, or even entire organisms. Multiple advanced aging biomarkers based on omics analyses and machine learning techniques, most notably epigenetic aging clocks but also other aging biomarkers, lend themselves for studying this process in greater detail. These biomarkers have been shown to predict the biological age of mammalian species including humans, mice, and naked mole rats, as well as many other species, and have become invaluable tools for tracking the aging trajectories during various biological processes^10–22^.

While most of the experimental approaches to rejuvenation have focused on cellular reprogramming, the application of various epigenetic clocks to mouse and human embryonic development led to a remarkable finding that rejuvenation may also occur naturally, reverting cells to a younger state^23^, approximately around the stage of mammalian gastrulation^24^. These studies offered the first evidence of a “ground zero” point of aging, where embryonic rejuvenation ends and organismal aging begins. However, the previous work had insufficient temporal resolution during embryonic development, with subsets of embryonic stages covered by data from multiple labs in separate experiments. It is also unclear whether the ground zero concept applies only to mammals or to a broader group of organisms. Therefore, further understanding of natural rejuvenation and associated mechanisms is needed in various animal species. A key obstacle to further advances in this area is that common mammalian animal models are poorly suited for such studies. Their embryos develop largely *in utero,* complicating longitudinal monitoring, and embryos are produced in small numbers and have a small size.

In contrast, the African clawed frog *Xenopus laevis* provides several advantages for the study of embryonic development^25^: 1) more than one thousand eggs can be produced at a time by a single female, 2) frog embryos are large (~1.2 mm diameter) and can be easily tracked during development, 3) their synchronous development is well characterized, and 4) *Xenopus* exhibits CpG methylation patterns that show difference with age and embryonic development^26^.

Here, we investigated changes in DNA methylation profiles, gene expression and incidence of developmental disruption throughout the embryonic development of *Xenopus laevis.* Using methylation microarrays, we found a decrease in DNA methylation entropy and an increase in embryo transcript abundance during early embryonic development. Based on skin DNA methylomes of adult *Xenopus laevis* varying in age, we developed the first frog epigenetic aging clock, which demonstrated epigenetic aging in this species. By applying this clock to *Xenopus* embryonic development, we discovered that epigenetic age declines approximately at the onset of gastrulation, pointing to a natural rejuvenation event. Interestingly, we also observed that embryos exhibit most developmental abnormalities during gastrulation. Taken together, our data quantify and advance the notion of ground zero of aging, a critical moment in vertebrate development, rejuvenation and aging.

## Results

### Profiling DNA methylation during aging and embryonic development in *Xenopus*

To characterize the dynamics of aging and development of the African clawed frog *Xenopus laevis,* we designed experiments to examine a set of aging parameters: epigenetic age, epigenetic entropy, global DNA methylation levels, global transcript abundance, and mortality due to disrupted development. To characterize age-related DNA methylation dynamics, we collected embryo samples during early development (**Fig. 1a**), spanning the period from 5 to 31.5 hours post-fertilization (hpf). In parallel, we continuously monitored and recorded a separate group of embryos from the same cohort to visualize progression through developmental stages. Under our experimental conditions, embryos began gastrulation at ~9.5 hpf and completed it 3 hours later. We also collected skin biopsies of adult animals throughout their lifespan (54 samples of *Xenopus laevis* aged 2-19 years old), isolated DNA from these samples and subjected them to DNA methylation profiling (**Fig. 1a**).

**Fig. 1.**
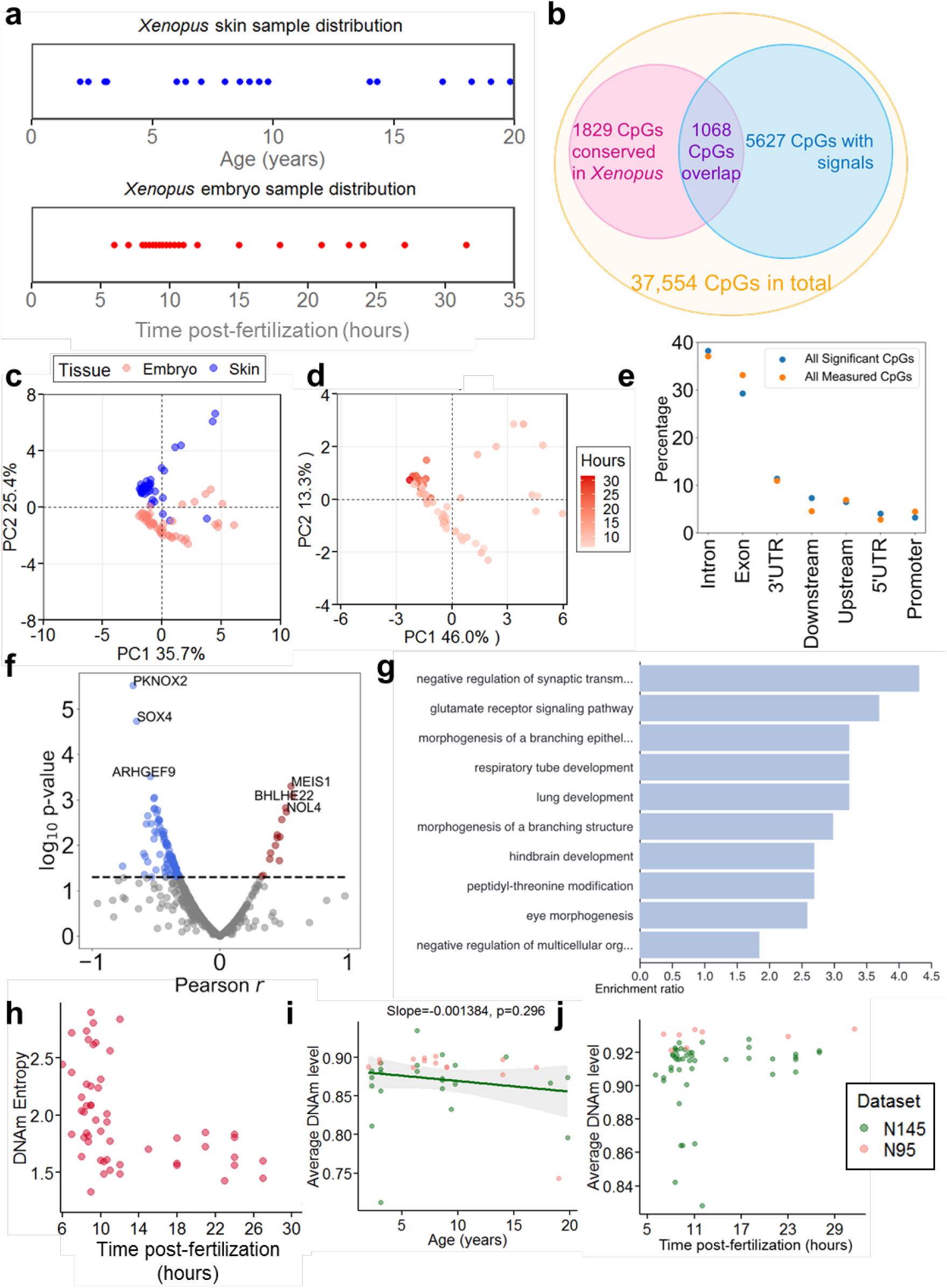
Methylation average and methylation entropy of *Xenopus laevis* samples. **a.** Age distribution of skin and embryo samples analyzed in the study. Skin samples were collected from animals aged 2-19 years, and embryo samples were collected from 6 and 31.5 hpf. **b.** Venn diagram showing all microarray sites, sites that showed significant signals, sites shared between microarray and the *Xenopus*-conserved CpGs by EWAS, and overlapping sites of both. **c.** Principal Component Analysis on all samples passing the QC based on significant sites overlapping with the *Xenopus*-conserved CpG sites. Each dot represents a sample, and tissue types are separated by colors. **d.** Principal Component Analysis of embryo samples passing the QC based on sites with significant signal overlapping with the *Xenopus*-conserved CpG sites. Deeper color indicates longer time post-fertilization. **e.** Percentage of all 1068 and 123 significant age-related genes in different gene regions. Intergenic upstream and downstream regions are shown as “upstream” and “downstream” **f.** Log_10_ p-value of the genes increased or decreased with age plotted as a function of Pearson correlation r with age. The 3 most significant genes with increased or decreased methylation are labeled. **g.** Enrichment analysis of genes with age-dependent gain or loss of DNA methylation. Enrichment levels for the top 10 gene ontology enriched biological processes are reported. **h.** DNA methylation Shannon entropy plotted as a function of development time based on sites with significant signal overlapping with the *Xenopus*-conserved CpG sites. **i-j.** Average DNA methylation level plotted as a function of skin age (**i**) and as a function of hours post-fertilization of embryos (**j**) based on the sites with significant signal overlap with the *Xenopus*-conserved CpG sites. Linear regression was performed to visualize the trend of entropy changes with skin age. N145 and N95 are two datasets that we collected, both spanning embryogenesis and adult ages.

*Xenopus* skin samples and embryos were analyzed by using the mammalian methylation array that was designed to profile highly conserved CpGs (**Fig. 1b**)^27^. From the samples we used, we generated 2 datasets, namely N95 and N145. We first found a total of 5,627 CpGs with significant values based on a hybridization signal. Although the microarray was designed to profile the methylome across mammals, we found in our EWAS study that 1,829 CpG sites were conserved in frogs, suggesting that the array may be useful for assessing methylation levels of many CpG sites in the *Xenopus* genome. By intersecting probes with the reliable signals and CpG sites conserved in *Xenopus*, we identified 1,068 CpG sites for further analyses. Of these CpGs, 109 sites significantly decreased DNAm levels with adult age, and 14 CpGs increased it (**Extended Data Fig. 1a**). To ensure quality of the data, we implemented a procedure (see Methods) to eliminate samples of low quality, filtering out 34 embryo and 16 skin samples (**Extended Data Fig. 1b-g**). These samples were evenly distributed across ages and were not biased toward a particular developmental stage (**Extended Data Fig. 1h**). Principal component analysis (PCA) showed that the samples clustered by tissue origin, with most skin samples having positive values and embryonic samples with negative values in PC2 (**Fig. 1c**). In addition, we carried out separate PCA of each tissue type and examined aging trajectories (**Fig. 1d, Extended Data Fig. 2a**). Embryo samples aligned with age diagonally in PC1 and PC2. Interestingly, older samples tended to be outliers in the skin PCA, whereas outliers in the embryo PCA were usually younger samples. This suggests that the aging trajectory of the embryos may differ from that of the aging skin.

We further examined CpG sites that significantly changes their methylation levels with age by mapping them onto the human genome (**Extended Data Fig. 1a**). We first analyzed the regions where these CpGs reside (**Fig. 1e**). CpGs significantly methylated or demethylated with age showed higher abundance in exons and lower in intergenic locations (however, note that the profiled CpGs were preselected based on conservation and therefore may be biased towards exons). We then plotted a volcano plot showing the genes highly correlated with age and their p values (**Fig. 1f**). Although we focused on all genes with age-related changes in methylation levels, many genes that regulate developmental processes were uncovered. For example, SOX4, a gene that was shown to regulate both aging and development^28,29^, appeared among the genes decreased most significantly with age (**Extended Data Fig. 2b**), and MEIS1, a transcription factor that regulates the developmental process and transcription in embryonic stem cells (**Extended Data Fig. 2c-d**), increased most significantly^30^. We additionally performed an enrichment analysis for the set of genes containing one or more sites whose DNA methylation levels were significantly increased or decreased with age (**Fig. 1g**). The enriched were developmental processes (respiratory tube, lung, and hindbrain development). Also, as previous findings suggested that age-dependent hypermethylation at polycomb repressive complex PRC2 regions is conserved in humans and mice^31,32^, our results suggest that such a mechanism may be conserved in frogs as well (**Extended Data Fig. 2e**).

Previous studies found an accelerated increase in DNA methylation Shannon entropy with adult age^21,33,34^, which is in line with the notion of accumulation of age-related damage. Using skin and embryo DNA methylation data, we examined the dynamics of methylation entropy in the frog. Methylation entropy of the skin had an increasing trend with age, although it did not reach significance (**Extended Data Fig. 2f**). In contrast, it decreased during early embryogenesis, reaching the lowest point at 10-12 hpf (**Fig. 1h**). We further separated the sites with and without significant methylation changes with age, and calculated entropy separately for them (**Extended Data Fig. 2g-j**). There was no difference between the entropy trends as a function of the embryonic age. However, skin DNA methylation entropy based on significant CpG sites increased more sharply with age (**Extended Data Fig. 2g-h**).

To examine the dynamics of global DNA methylation during aging and embryonic development, we calculated average global DNA methylation (**Fig. 1i-j**). Skin showed a slight decreasing trend with age, whereas embryonic methylation had an increasing trend in early development, reaching a plateau around 10 hpf. Taken together, our data suggest that global DNA methylation reaches a plateau and achieves the lowest entropy state approximately at the gastrulation stage of embryonic development.

### Gene expression dynamics during *Xenopus* embryonic development

In addition to the collected DNA methylation, we examined publicly available transcriptomic data derived from eggs through embryonic stage NF35^35^ (**Fig. 2a**). To allow for data cross-comparison, we transformed embryonic stages to the approximate time post-fertilization and gastrulation. To examine changes in gene expression patterns during embryonic development, we utilized adult *Xenopus* organ samples to identify highly expressed genes, and calculated their gene expression levels across development. PCA showed that PC1 decreased with embryonic development, whereas PC2 increased during early development and then decreased (**Fig. 2b**). To determine the exact inflection point of these two principal components, we plotted them against time post-fertilization (**Fig. 2c-d, Extended Data Fig. 3a-b**). Interestingly, PC1 had the highest value just before the beginning of gastrulation (NF8-9) and then sharply decreased. This trend then flattened with time. PC2 had the highest peak that occurred right after gastrulation begins, then the value decreased. We then calculated mean counts per million (CPM) gene expression levels as a function of embryonic age (**Fig. 2e, Extended Data Fig. 3c**). We found that the lowest expression was at embryonic stage NF9 (just before gastrulation), followed by a dramatic increase during gastrulation. After the gastrulation stage, the increasing trend gradually flattened.

**Fig. 2.**
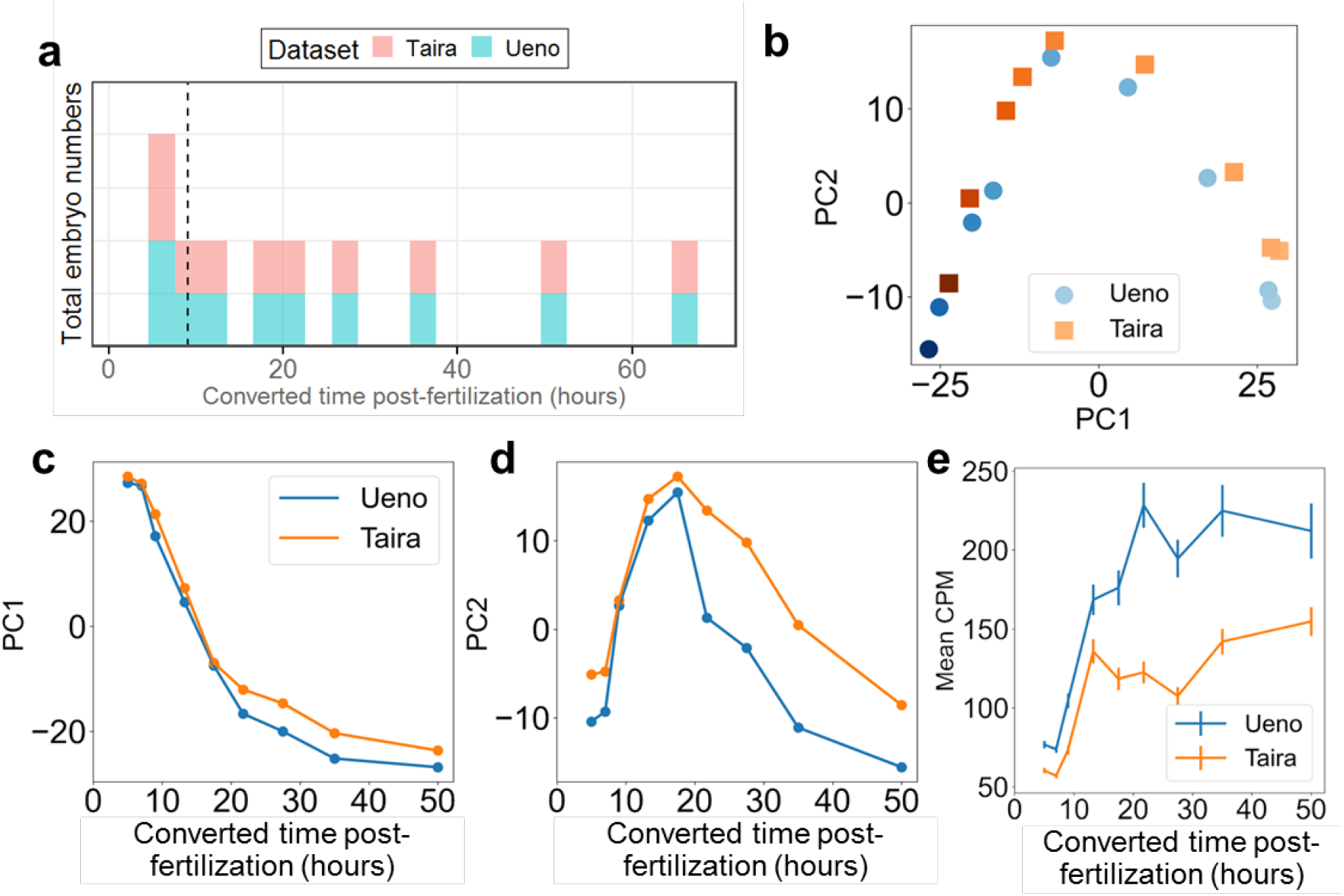
Global transcription of *Xenopus* embryo samples. **a.** Number of samples used in each dataset as a function of hours post-fertilization converted from embryonic stages. Gastrulation starting time is marked with vertical dash lines. **b.** Principal component analysis of gene expression levels of *Xenopus* embryos. Datasets are separated by colors. **c-d.** Principal Component analysis of transcription data. PC1 (**c**) and PC2 (**d**) are plotted as a function of embryonic age in hours. **e**. Mean CPM (counts per million) gene expression levels as a function of embryonic age in hours.

### Development of the *Xenopus* epigenetic aging clock

Our recent studies revealed a decrease in epigenetic age during early embryogenesis in mice followed by an increase starting approximately at gastrulation, with the minimal epigenetic age corresponding to ground zero of aging^24^. Such analyses in *Xenopus* embryos could not be done, as epigenetic aging clocks do not exist for this species or for any amphibian. To develop such a clock, we employed a bootstrap aggregated ElasticNet model and trained an epigenetic aging clock based on adult *Xenopus* skin samples ages 2-19 years (**Fig. 3a-e**). To identify the optimal hyperparameter (Lambda=0.0039) and the subset of 185 optimal clock CpGs, we used out-of-sample cross-validation applied to the whole 40 samples. Then, we used 40 adult skin samples to bootstrap 40 out-of-sample ElasticNet models, in each of which 1 sample was used as a test set, and the remaining 39 samples were used as a training set. The quality of predictions was estimated based on the test samples, and then averaged over all 40 bootstrapped models. The resulting clock was the average of predictions of all bootstrapped models, and showed MAE of 1.82 years on the adult samples not used in the training.

**Fig. 3.**
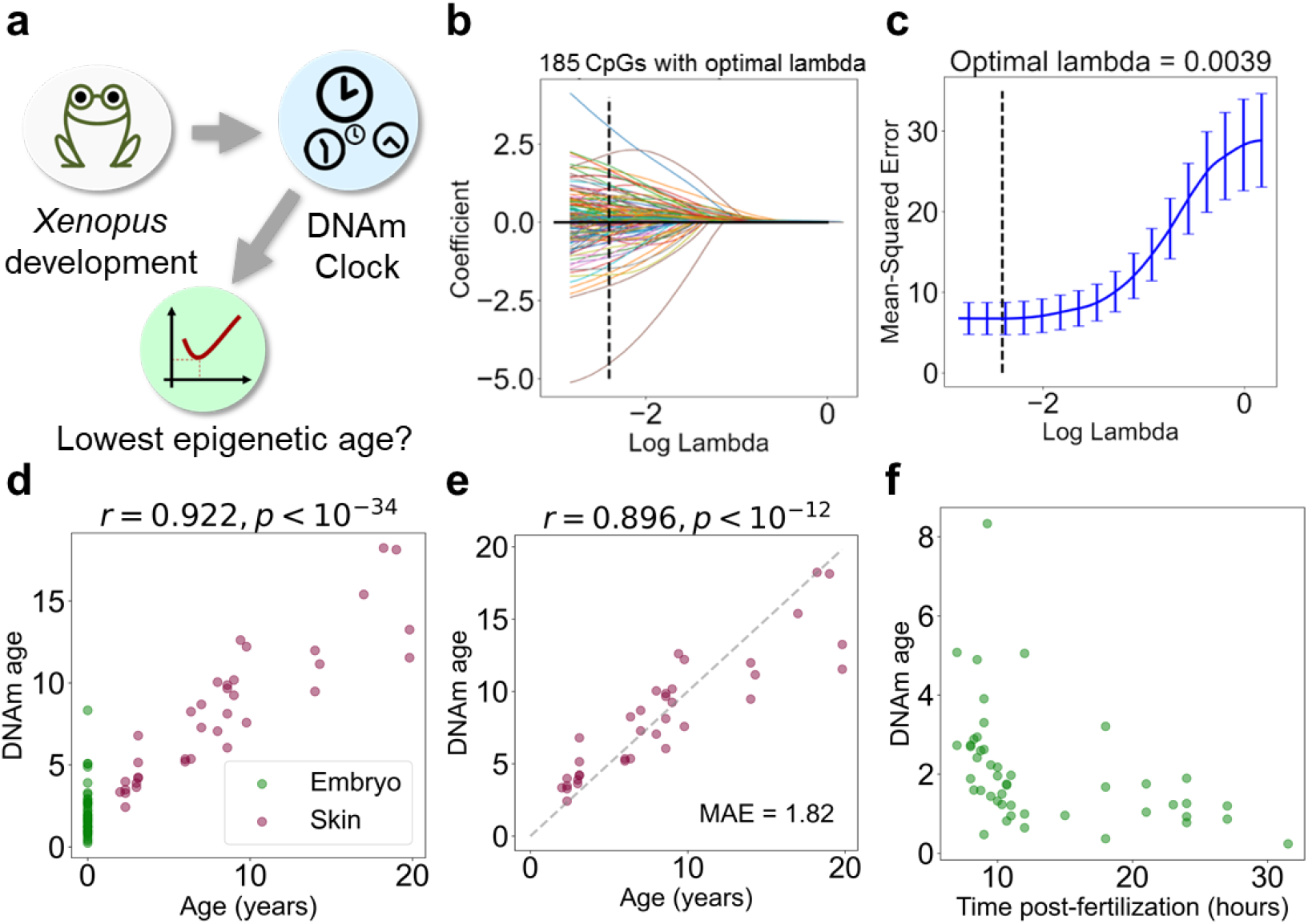
Development of the frog epigenetic aging clock and its application to *Xenopus* embryo samples. **a.** Schematic of experimental and data analysis settings. **b.** Out-of-sample cross validation path for ElasticNet regression trained on 40 adult skin samples. The optimal number of CpGs is 185. **c**. Mean-squared error cross validation path for ElasticNet regression. The optimal lambda is 0.0039. **d**. Bagged out-of-sample trained ElasticNet regression trained on adult skin samples and applied both to embryo and skin samples. **e**. Bagged out-of-sample ElasticNet regression trained on skin samples. **f.** DNAm age prediction of the clock trained on adult skin samples and applied to embryo samples.

The data clearly show that *Xenopus laevis* epigenetically ages during adult ages, a feature that was preivously described for all tested mammals regardless of their mortality patterns. Moreover, application of the *Xenopus* skin aging clock to embryonic samples revelaed their predicted low biological age (**Fig. 3d**), suggesting that this clock also captures the biological age of embryos.

### *Xenopus* epigenetic aging clock reveals the lowest epigenetic age during gastrulation

By applying our epigenetic aging clock to frog samples throughout embryonic development, we observed a somewhat similar pattern of epigenetic age dynamics as in case of DNA methylation entropy (**Fig. 3f**), i.e. the epigenetic age declined rapidly around 10-12 hpf, at the onset of gastrulation, where it reached its minimum, then very slowly increased. Notably, the high epigenetic age at early stages is not due to the low-quality samples (**Extended Data Fig. 4a-f**), and we additionally found significant correlations between DNA methylation average level, entropy and epigenetic age (**Extended Data Fig. 4g-h**). These data are consistent with our previous analyses in developing mouse embryos that showed that the epigenetic age decreases during early embryogenesis, reaching its minimal point around gastrulation, then begins to increase^24^. Thus, the data suggest that the *Xenopus* embryos are also rejuvenated during early embryogenesis and then begin to age approximately at the stage of gastrulation, establishing aging ground zero in this species.

### Increased incidence of developmental disruption during *Xenopus* gastrulation

Gastrulation corresponds to a critical period of organismal life, which is associated with the formation of germ layers, establishment of body plan and separation of the germline and soma^36,37^. To test how this period is related to mortality, we carried out a large-scale analysis, following the developmental trajectories of 6,457 embryos (**Fig. 4a**). For this, we developed an approach wherein eggs were fertilized *in vitro* and loaded individually into separate wells of 384-well plates. The vast majority of these embryos were scanned from around the 8-cell stage for at least 24 hours. We collected images every 15 minutes, generating videos and using them to determine the timing of embryonic developmental stages and death. We aggregated the data from a total of 7 trials, with 240-1904 embryos each. To ensure that the eggs were fertilized, we excluded those that did not fertilize or symmetrically cleave, starting the experiment from 4-cell embryos. This approach excluded over 1,000 embryos due to issues related to fertilization, initial asymmetric cleavage, stalling after the first cleavage and pseudo-cleavage. The absolute majority of the remaining 5,546 embryos developed normally until at least stage 22, i.e. they showed no abnormal development for ~24 hours post-fertilization. However, among these 5,546 embryos, 104 embryos exhibited disrupted development within 24 hours of fertilization and died (**Fig. 4b**). Some common features found in the embryos that exhibited such severe developmental abnormalities were that their cells had abnormal asymmetrical division just pre-gastrulation, mis-formation of the blastopore ring in early gastrulation, and failure of blastopore contraction during gastrulation (**Fig. 4c**). These severe abnormalities typically led to embryonic developmental interruption. To more accurately pinpoint the embryonic stage and developmental time at which these severe developmental abnormalities occurred, we chronologically ordered embryonic developmental disruption events. More than half of these 104 embryos had developmental interruptions that occurred around the onset of gastrulation, immediately often following blastopore ring formation. Since the time interval between fertilization and gastrulation in different batches of embryos was slightly different, we set time zero as the time when we first observed blastopore ring fragments (**Fig. 4d**). This normalization allowed us to overcome the effects of slight variation in temperatures and batches, thereby enabling comparison across experiments. These normalized data showed that 66 embryos exhibited critical abnormal developmental manifestations within the first 3 hours following the onset of gastrulation, indicating that this developmental stage features disproportionately high mortality in embryonic development. Nevertheless, it should be stressed that the overall embryonic mortality after the 4-cell stage was quite low, around 2%.

**Fig. 4.**
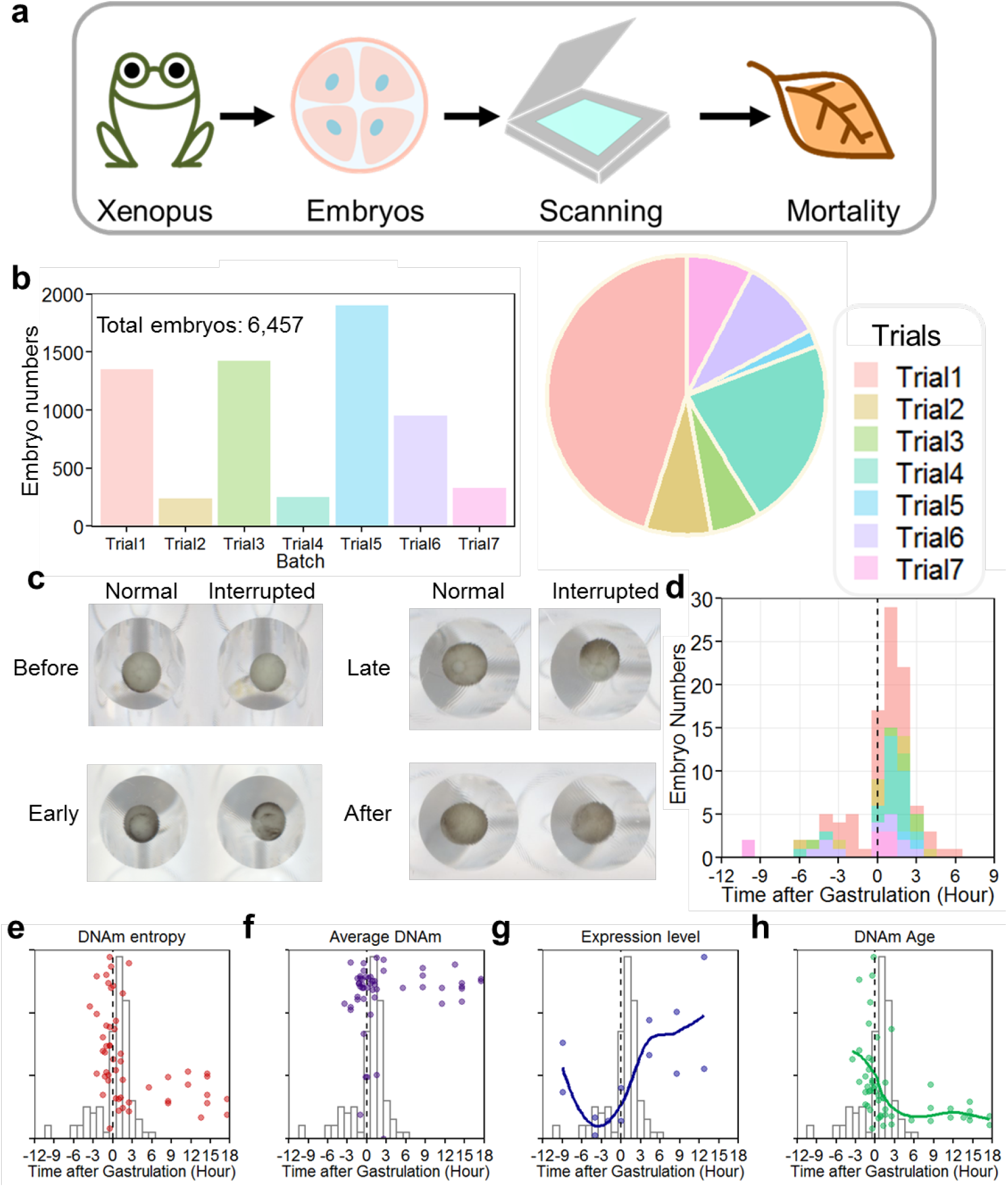
Analyses of embryonic mortality and its association with predicted biological age and other aging parameters. **a.** Schematic of the experiment. *Xenopus laevis* eggs were collected, fertilized and loaded into 384-well plates, then scanned in the form of time-series images to monitor develomental disruption and mortality. **b.** Number of samples used in each trial analyzed by the mortality assay. **c.** Pie chart containing the percentage of embryos that exhibited developmental interruption. Different batches are separated by color. Total number of embryos is 104. **c.** Representative scans of embryos that stopped development before gastrulation, during early gastrulation, during late gastrulation and after gastrulation with normally developed embryos from the same batch. Videos can be found in Supplementary Information. **d**. Numbers of embryos that exhibited developmental interruption plotted as a function of embryonic time. Time normalized by setting the starting point of gastrulation as point zero. Different batches are separated by color. **e-h.** Co-plotting of normalized DNA methylation Shannon entropy (**e**), global DNA methylation level (**f**), global gene expression (**g**) and epigenetic age (**h**) with the number of embryos that stopped development, plotted as a function of embryonic time after gastrulation. Time normalized by setting the starting point of gastrulation as point zero. For gene expression, embryonic time was converted from embryonic stages.

We further characterized the relationship between the point of maximum mortality and other aging-related patterns we observed above (i.e. epigenetic age, entropy, global transcript levels, global DNA methylation) (**Fig. 4e-h**). The lowest DNA methylation entropy and epigenetic age, as well as the highest global DNA methylation corresponded to the highest developmental disruption/mortality rate. This stage also featured a rapid increase in embryo transcripts. All these features map to gastrulation, pointing to ground zero of aging during this stage of frog development.

## Discussion

When does aging of an organism begin? Arguably, this point should correspond to the lowest biological age of that organism^38^. The germline is composed of metabolically active cells, which accumulate damage over time, thereby becoming older. If we neglect the negative selection of unviable embryos, the absence of a rejuvenation event would lead to the age of the zygote above zero, and its increase through generations. Such a situation would be unsustainable for the survival of a species. Therefore, for each generation to begin aging at the same lowest biological age, there is likely a rejuvenation event during embryonic development. Such an event was recently described in mammals, with the experiments carried out primarily in mice. Its exact timing could not be pinpointed, but it corresponded approximately to the period of E4.5-E9.5, when the minimal biological age was reached. We have termed this point the ground zero of aging^24^.

Prior to our current study, it was unclear whether this natural rejuvenation event and the ground zero timepoint were limited to mammals or also occurred in other groups of organisms. Also, comprehensive datasets with high temporal resolution describing methylation dynamics during the entire developmental process are lacking in any species, precluding localization of the exact timing of these events. Such studies in mammals are difficult as embryos cannot be readily observed or studied in large numbers. These limitations may be avoided in the *Xenopus laevis* model, where oocytes are large (more than 1 mm) and can be conveniently fertilized *in vitro* in large numbers and then monitored, imaged, or collected for molecular analyses.

One limitation of the *Xenopus* model, however, is the lack of epigenetic aging clocks that could be used to assess changes in biological age. To address this deficiency, we took advantage of our aging *Xenopus* colony at Harvard Medical School and collected skin biopsies from 2-19-year-old frogs. We found that the mammal methylation array can be used to study DNA methylation in *Xenopus laevis*, although the number of usable CpG sites was limited to about one thousand. With this tool, we were able to develop the first epigenetic aging clock of *Xenopus laevis*, with MAE of less than 2 years. Application of this clock to embryonic samples revelaed their low biological age. Thus, frogs epigenetically age, like all previously examined mammals, and the skin clock we developed could be used to assess the age of embryos.

Application of our frog clock to samples throughout embryonic development revealed a rapid rejuvenation event during early embryogenesis, wherein the minimal age was reached around 10 hpf, which corresponds to the gastrulation stage. This observation suggests that, like mammals, frogs are epigenetically rejuvenated around gastrulation and that their biological aging begins afterwards. We further found that the rejuvenation event corresponds to the lowest DNA methylation entropy, and a sharp increase in embryonic global transcript abundance, pointing to the critical importance of that stage of development for aging and rejuvenation.

This stage involves the establishment of body plan, separation of the germline and soma and active embryonic transcription. These most fundamental processes may be prone to errors, due to their complexity and contribution of genetic burden. To test this possibility, we examined embryonic mortality of the frogs by simultaneously monitoring developmental trajectories of thousands of embryos. This method involved *in vitro* fertilization of eggs, followed by the distribution of individual embryos into 384-well plates and monitoring and recording the plates over the period of 24 hours. Embryonic mortality is thought to be high in mammals, where it was suggested that more than 50% of conceptions do not reach the stage of puberty, consistent with the K-reproductive strategy minimizing investments into unviable progeny at the earliest stages of development. However, detailed quantification of mortality in mammals or other vertebrates is lacking. In our study, of more than five thousand 4-cell stage embryos analyzed, only 104 died. Thus, consistent with the r-reproductive strategy delaying negative selection of unviable embryos to later stages of life, frog embryos exhibit low embryonic mortality after the 4-cell stage, on the order of ~2%. However, more than half of the embryos with disrupted development showed defects and/or died during gastrulation, strongly supporting the model of aging ground zero during this stage of development. It should be noted, however, that the assay we employed defines mortality based on the timing when embryos display crucial developmental deficits and it may be restricted by the limitations of scanning resolution. Future work based on high-throughput high-resolution microscopy may improve the analyses of developmental mortality.

It would be interesting to extend the findings with additional embryos as well as develop and apply advanced aging biomarkers (preferably multi-tissue and multi-omic). Additional opportunities arising from our study are that it established an experimental system for mechanistic analyses of embryonic rejuvenation. As such, it may help characterize commonalities and differences in embryonic rejuvenation between amphibians, mammals and other vertebrates.

Overall, our study suggests that frogs, like mammals, epigenetically age and that they exhibit a rejuvenation event during early embryogenesis, culminating in ground zero at the stage of gastrulation, which defines the beginning of their aging. Lewis Wolpert once famously said, “It is not birth, marriage, or death, but gastrulation which is truly the most important time in your life.”^39,40^ It may now be said that this stage is also the most important time for aging and rejuvenation.

## Acknowledgements

Supported by the Impetus grant program, the Michael Antonov Foundation and NIA grants to VNG. LP and WR were supported by the Paul G. Allen Frontiers Group and McKenzie Family Charitable Trust.

## Methods

### Animal experiments

All *Xenopus laevis* animals were from the Harvard Medical School facility, and all animal protocols were approved by the local IACUC. We aged frogs locally as well as acquired the oldest frogs over the years from the Marine Biological Laboratory and Cold Spring Harbor Laboratory. Frogs were kept in tanks with constant temperature (18 °C) and 12/12 hour light cycle. We collected skin biopsies from frogs of different ages (age and gender of the samples are shown in **Supplementary Table 1**). Oocyte collection from females, *in vitro* fertilization and de-jelly steps followed published protocols^41^. The embryos were then kept in 1x MMR buffer, sorted at the 4-cell stage to remove unfertilized and abnormal embryos, and then individually loaded into 384-well plates (Phenix, MPU-8210) pre-loaded with 70 ul 1x MMR buffer. The plates were scanned every 15 min throughout the next 24 hours. In parallel, embryos were collected for molecular analyses. We combined different numbers of samples for these analyses to approximately match the total DNA in the samples (i.e. with the development stage, the number of samples of pools decreased).

### Isolation of genomic DNA

DNA was isolated from *Xenopus laevis* skin samples and embryos using the DNeasy Blood and Tissue Kit (Qiagen) following the manufacturer’s protocol. Concentration of DNA samples was determined using the Qubit dsDNA BR assay kit (Invitrogen). Isolated DNA was stored at −80°C before being applied to microarrays.

### DNA methylation profiling

Two batches of samples were applied to DNA methylation profiling, generating 2 datasets respectively named N95 and N145. Details about the two datasets shown in **Extended Data Fig. 1g**. Methylation data was generated through the Epigenetic Clock Development Foundation, subjected to the HorvathMammalMethylChip40^27^. Samples were randomized to avoid introduction of batch/chip position effects. All sample preparation/processing was carried out according to the Illumina kit protocols. For microarray analysis, raw methylation data were first normalized using the SeSAMe R package and beta values were calculated. To calculate Shannon entropy, the DNAm levels’ probability density distribution for each sample was discretized into 30 bins and calculated, and later used as an input to scipy.stats.entropy Python 3.9 package. Human Genome Reference Consortium GRChg38 was used for gene annotations. Enrichments analysis of genes was performed using WebGestalt 2019 ORA based on the gene ontology biological process database^42^. Normalized PRC2 binding levels were calculated based on average EZH2 and SUZ12 Chip-Seq binding levels from mouse embryonic cells lines (E14) publicly available under accession numbers GSM2472741 and GSM3243624^32,43^.

### *Xenopus laevis* genome annotation

We mapped the probe sequences of the mammalian methylation array to the reference genome Xenopus_laevis 10.1 from ENSEMBL. The alignment was done using the QUASR package, which accounts for the bisulfite conversion treatment. The CpGs were annotated based on the distance to the closest transcriptional start site using the ChIPseeker package^44^.

### Sample CpG filtering

We adapted the HorvathMammalMethylChip40 to amphibian *Xenopus laevis* DNA sequences. To preselect evolutionary conserved CpGs, we used the following method. We identified 1,829 CpG sites present in the array that were conserved in *Xenopus laevis*. By further analysis of the DNAm chip readout, we filtered out the hybridization of CpG sites that did not meet the quality control due to low signal’s intensity wherein the readout of methylation was set close to 0.5 due to technical realization of the chip. For 761 out of 1,829 CpGs, hybridization was insufficient, thus we filtered them out, and focused on 1,068 CpGs which were measured in most samples (**Extended Data Fig. 1**). Specifically, the filtering required that the average methylation across all samples deviated from 0.5 by at least 0.1, and the standard deviation of methylation changes across the samples was larger than 0.5. Additionally, we filtered out 43 samples where 1,068 hybridized frog CpGs were not properly measured (**Extended Data Fig. 1**). In total, out of 142 samples and 37,554 CpGs, 99 samples and 1,068 CpGs passed the initial quality control.

An additional quality control score for DNAm clock bootstrap aggregated ElasticNet predictions was calculated as the fraction of clock CpGs sites (185 CpGs) that were measured by the chip with a significance level higher than p<0.05. The exclusion threshold was chosen at the level of 55% clock CpGs measured in a sample. Out of 99 samples, 7 failed the second quality control and were excluded from the analysis: 2 adult skin samples, and 5 embryo samples. It is worth noting that measured frog methylation levels were strongly biased towards hypermethylated regions.

### DNAm clock analysis

For training the DNAm clock, we employed a bagged ElasticNet model trained with out-of-sample cross validation. As the training set, we used 40 adult skin samples. The optimal hyperparameter lambda = 0.0039 for ElasticNet was identified by out-of-sample cross validation, as well as the optimal number of CpGs — 185 out of 1,068. For prediction, we trained 40 models, and as a predictor we employed the bagged ensemble of these 40 models. We also applied the bagged ElasticNet to embryonic samples for prediction of their epigenetic age.

### Transcriptomic data analysis

RNA-seq data were obtained from Taira (2012.03) and Ueno (2012.10) datasets in GEO database (GSE73430) from^15^. Adult samples were used for normalization to determine the set of transcripts to be analyzed. Time post-fertilization and gastrulation were transformed from^45^. Principal components and RNA expression levels plotted as a function of the NF stage are shown in **Extended Data Fig. 3**.

### Embryonic mortality and developmental disruption assays

Embryos that did not reach the NF stage 3 (4-cell stage) were excluded from the analysis. Epson V600 perfection photo scanner was used to capture embryonic pictures in the form of time series. Scanning videos and a complete list of scanning intervals are shown in Supplementary Infomation. Embryos were further examined to remove non-developing embryos. The remaining embryos that did not complete development after 24 hours were scanned at ~22.2 °C. Embryos that developed a visible aberrant pattern or did not change morphology throughout time series were considered developmentally interrupted. A complete set of embryos analyzed is in Supplementary Infomation.

