## Extended Data Figures for "Epigenetic profiling and incidence of disrupted development point to gastrulation as aging ground zero in *Xenopus laevis*"

**Extended Data Fig. 1**

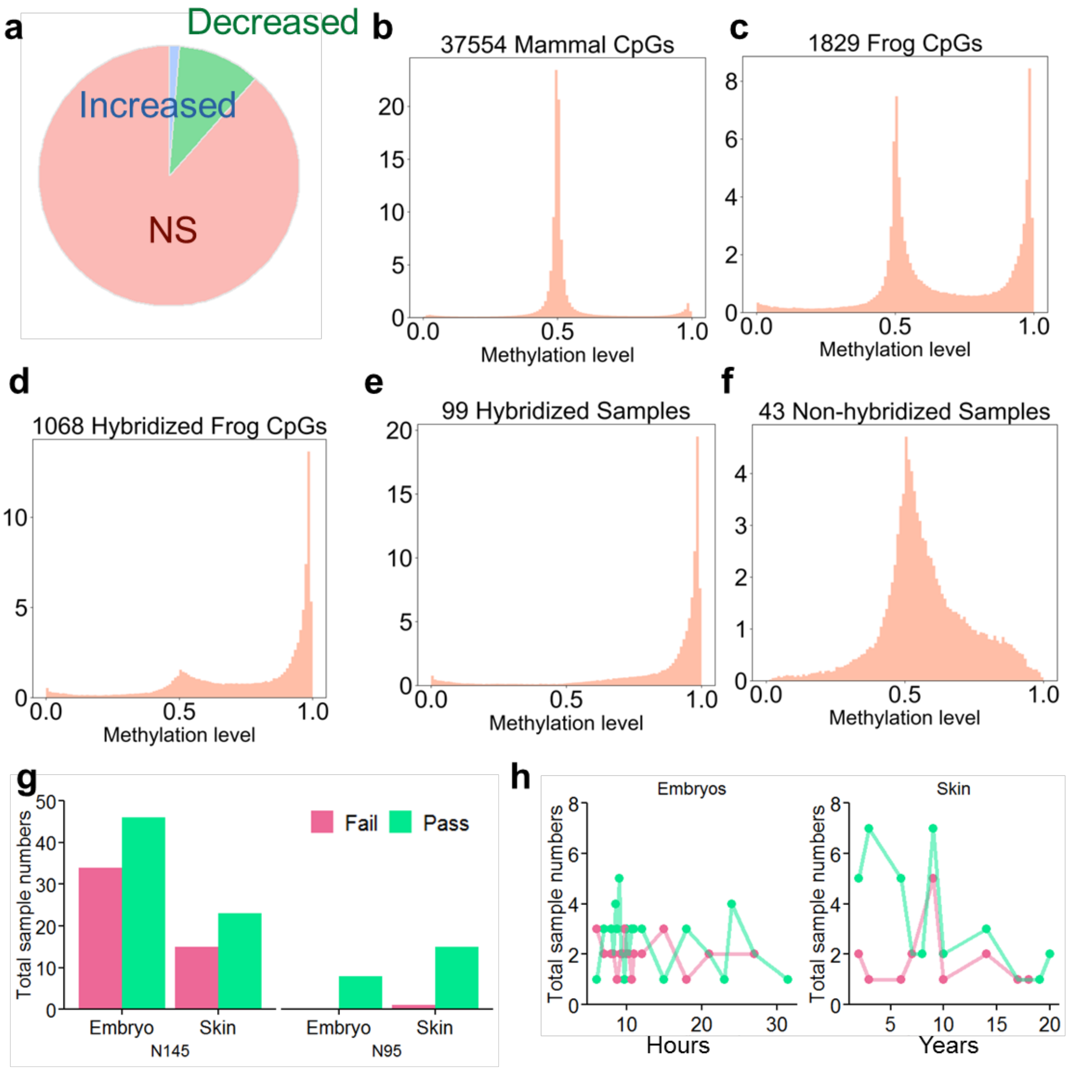


**Extended Data Fig. 1. a.** Percentage of hybridized significant CpG sites with significantly decreased (green), significantly increased (blue) and non-significant (pink) DNAm levels with age. **b-d.** Histogram of methylation levels across all samples measured (**b**) for all mammal CpG sites, (**c**) for frog CpG sites, (**d**) for hybridized CpG sites. **e-f.** Histogram of methylation levels across hybridized frog CpG sites for (**e**) hybridized samples, (**f**) non-hybridized samples. **g.** Number of samples that passed or failed QC in each dataset. Data are separated by tissue types. **h.** Number of samples that passed or failed QC as a function of time. Data are separated by tissue type.

**Extended Data Fig. 2**

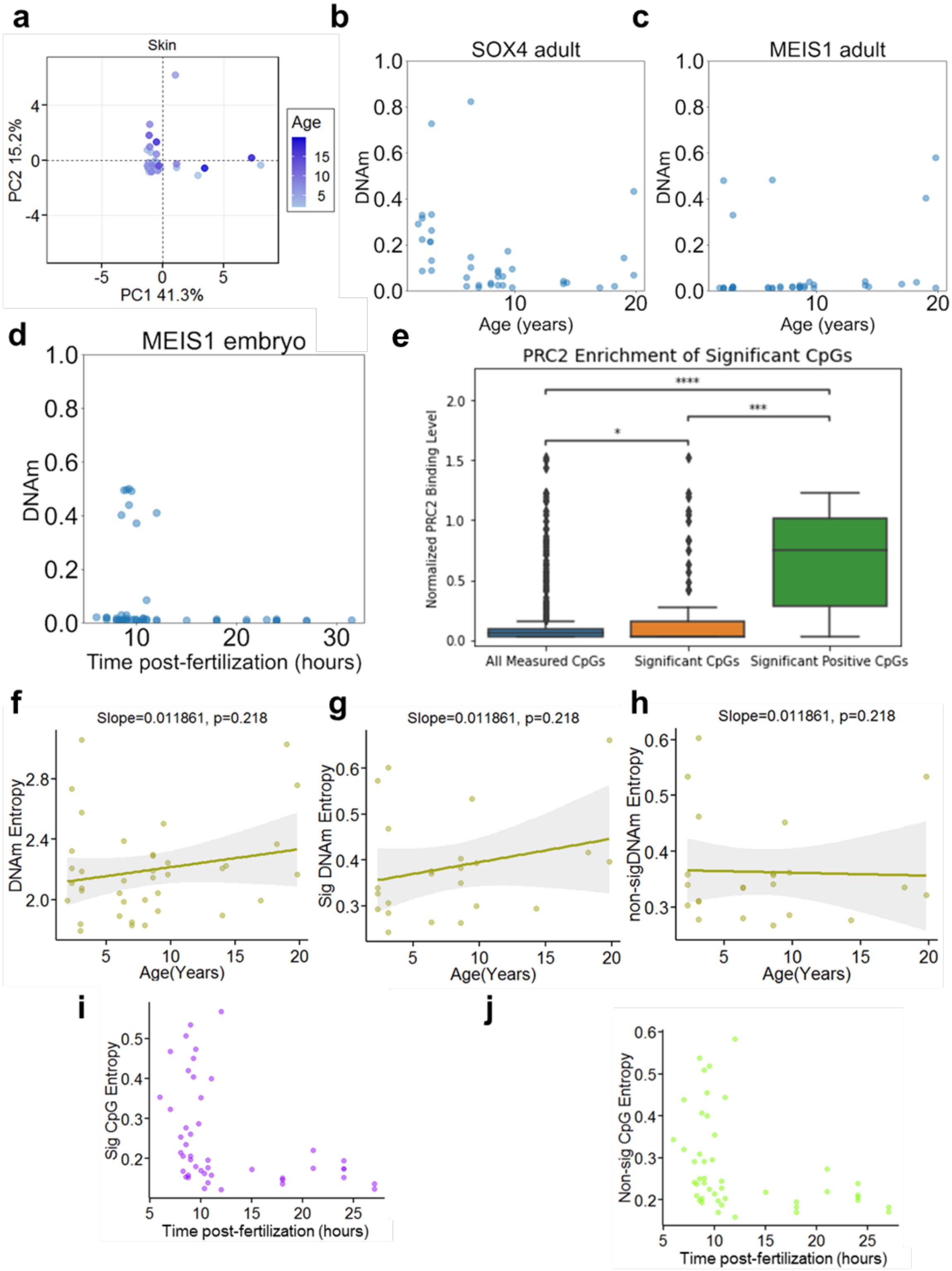

**Extended Data Fig. 2. a.** Principal Component Analysis of skin samples passing the QC based on significant sites overlapping with the *Xenopus*-conserved CpG sites. Age shown in years and deeper color indicates higher age. **b-c.** DNA methylation levels of CpG sites from genes SOX4 (**b**) and MEIS1 (**c**) plotted as a function of age in *Xenopus laevis* skin. **d.** DNA methylation levels of CpG sites from MEIS1 (**d**) plotted as a function of age in *Xenopus laevis* embryos. **e.** CpG sites that gain DNA methylation with age in the skin of frogs are enriched for PRC2 binding. p-value levels: * < 0.05, **: < 0.01, ***: < 0.001, ****: < 0.0001**. f-h.** DNA methylation entropy of all hybridized CpG sites (**f**), significantly decreasing hybridized CpG sites with age (**g**) and non-significant sites (**h**) plotted as a function of skin age. **i-j.** DNA methylation Shannon entropy plotted as a function of hours post-fertilization in embryo samples based on *Xenopus*-conserved CpG sites significantly decreasing with age (**i**) and not significantly changing with age (**j**).

**Extended Data Fig. 3**

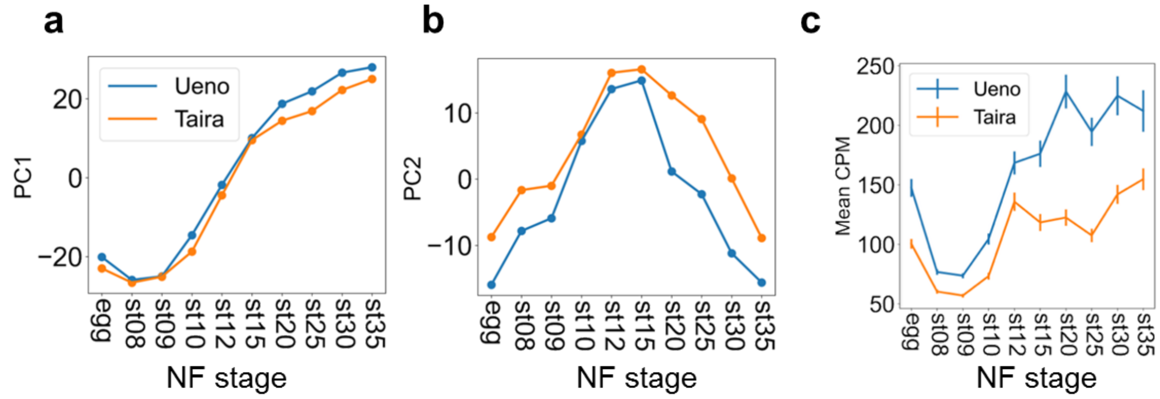


**Extended Data Fig. 3. a-c**. Principal component 1 (**a**), Principal component 2 (**b**), and gene expression levels (**c**) plotted as a function of the NF stage.

**Extended Data Fig. 4**

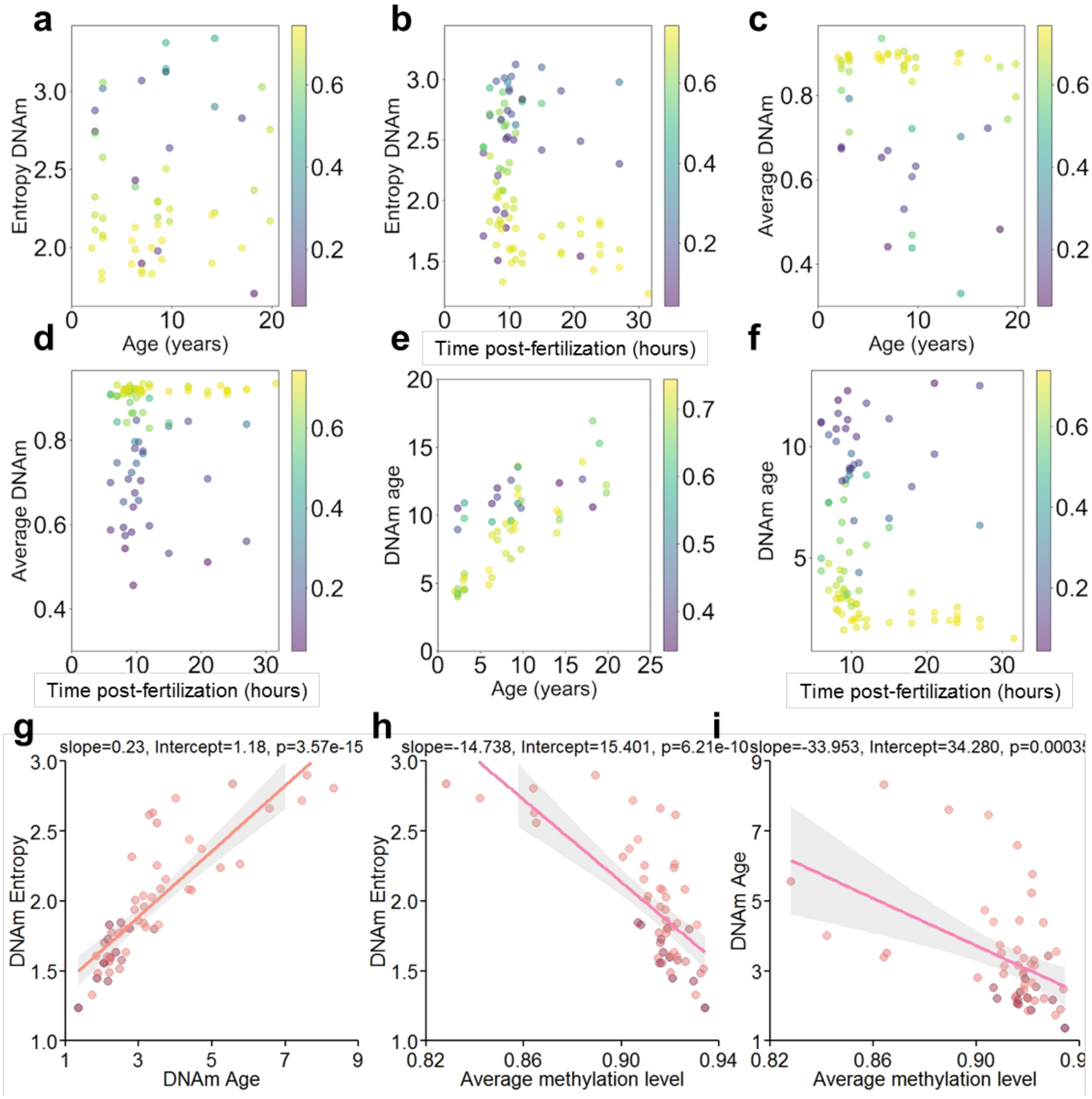


**Extended Data Fig. 4. a.** DNA methylation Shannon entropy of skin samples plotted as a function of development time based on the sites with a significant signal overlap with the *Xenopus*-conserved CpG sites. All skin samples analysed are shown, with the sample QC scores shown by color. **b.** DNA methylation Shannon entropy of *Xenopus laevis* embryo samples plotted as a function of development time based on the sites with a significant signal overlap with the *Xenopus*-conserved CpG sites. All embryo samples analysed are shown, with the sample QC scores shown by color. **c-d.** Average DNA methylation level plotted as a function of age of the skin (**c**) and as a function of hours post-fertilization in embryos (**d**) based on the sites with a significant signal overlap with the *Xenopus*-conserved CpG sites. All skin and embryo samples analysed are shown, with the sample QC scores shown by color. **e.** Bagged out-of-sample ElasticNet regression trained on all skin samples. All skin samples analysed are shown. Sample QC scores are shown by color. **f.** DNAm age prediction based on the clock trained on adult skin samples and applied to embryo samples. All embryo samples analysed are shown. Sample QC scores are shown by color. **g.** DNAm Shannon entropy plotted as a function of epigenetic age prediction. **h.** DNAm Shannon entropy plotted as a function of average DNA methylation. **i.** Epigenetic age plotted as a function of average DNA methylation.
